# Stability of navigation in catheter-based endovascular procedures

**DOI:** 10.1101/2023.06.02.543219

**Authors:** Chase M. Hartquist, Jin Vivian Lee, Michael Y. Qiu, Charles Suskin, Vinay Chandrasekaran, Halle R. Lowe, Mohamed A. Zayed, Joshua W. Osbun, Guy M. Genin

## Abstract

Endovascular procedures provide surgeons and other interventionalists with minimally invasive methods to treat vascular diseases by passing guidewires, catheters, sheaths and treatment devices into the vasculature to and navigate toward a treatment site. The efficiency of this navigation affects patient outcomes, but is frequently compromised by catheter “herniation”, in which the catheter-guidewire system bulges out from the intended endovascular pathway so that the interventionalist can no longer advance it. Here, we showed herniation to be a bifurcation phenomenon that can be predicted and controlled using mechanical characterizations of catheter-guidewire systems and patientspecific clinical imaging. We demonstrated our approach in laboratory models and, retrospectively, in patients who underwent procedures involving transradial neurovascular procedures with an endovascular pathway from the wrist, up in the arm, around the aortic arch, and into the neurovasculature. Our analyses identified a mathematical navigation stability criterion that predicted herniation in all of these settings. Results show that herniation can be predicted through bifurcation analysis, and provide a framework for selecting catheter-guidewire systems to avoid herniation in specific patient anatomy.

## 1 Main

Endovascular procedures have transformed the treatment of disease through by enabling surgeons and other interventionalists to advance endovascular tools to a distal vascular treatment site through a small vascular puncture site. The interventionalist typically first advances a guidewire through the vascular lumen, and then advances additional catheters and sheaths over this wire in a telescoping fashion. Once in place, tools for intervention are then advanced to facilitate the desired treatment. This minimally invasive approach is typically a safer and more efficient alternative to invasive open surgery [1]. However, endovascular procedures are plagued by a phenomenon known as herniation, where the catheter-guidewire system forms an extended loop and spontaneously shifts from the intended endovascular path, preventing the interventionalist from advancing the system to the desired distal vascular site. The interventionalist must often withdraw the catheter-guidewire system from the patient and repeat the attempt to reach the distal vascular site, not uncommonly with a completely new catheter-guidewire system. In part because the physics underlying herniation has not yet been elucidated, herniation cannot currently be predicted or prevented.

A range of new technologies are under development to circumvent the challenge of herniation entirely. Robotic navigation technologies under development may potentially enable interventionalists to bypass these problematic navigation procedures [2], and combinatory devices have emerged to reduce the need for multiple coaxial devices in surgery [3]. However, advanced implementations of these remain far from commercialization. Steerable devices offer improved navigational capabilities, which could potentially reduce herniation events [2, 4, 5], but these are also limited in practice. There thus continues to be a pressing need to understand and control herniation.

The impact of endovascular procedures and herniation are particularly evident in the management of patients with neurovascular conditions such as stroke, which impacts over 800,000 patients annually in the United States alone and is the primary cause of disability in the developed world [6]. In these procedures, endovascular tools are advanced toward the brain neurovasculature to restore blood perfusion, stop hemorrhage, or remove occlusive blood clots that lead to cerebral ischemia. The speed with which the blood perfusion is restored is the key determinant of patient outcomes. While endovascular procedures have transformed the treatment of acute ischemic stroke, the capacity to rapidly restore blood perfusion to the brain is highly dependent upon the efficiency of the catheter-guidewire system advancement without heniation [7–13].

The cost of herniation is also particularly evident in these procedures, with delays of only minutes leading to substantially worse patient outcomes [6]. To minimize treatment time, a vascular pathway commonly used in these surgeries is the transradial access (TRA) route, where the catheter-guidewire system is introduced to the vasculature through the radial artery (Fig. 1a) [14, 15]. However, as the catheter-guidewire system bends through the aortic arch, herniation can commonly occur. This can be seen in a sequence of fluoroscopy images acquired during a neurovascular procedure using TRA, with herniation occurring as a catheter was passed over a guidewire around the aortic arch (Supplementary Video 1). Although the interventionalist successfully tracked the guidewire through the aortic arch and up the carotid artery (Fig. 1b), the guidewire herniated down into the aortic arch as a catheter was advanced over the guidewire, forming an extended loop (Fig. 1c), requiring the surgeon to remove the catheter-guidewire system that failed and restart the procedure with a different, sterile catheter-guidewire system, which extended procedure time.

**Fig. 1.**
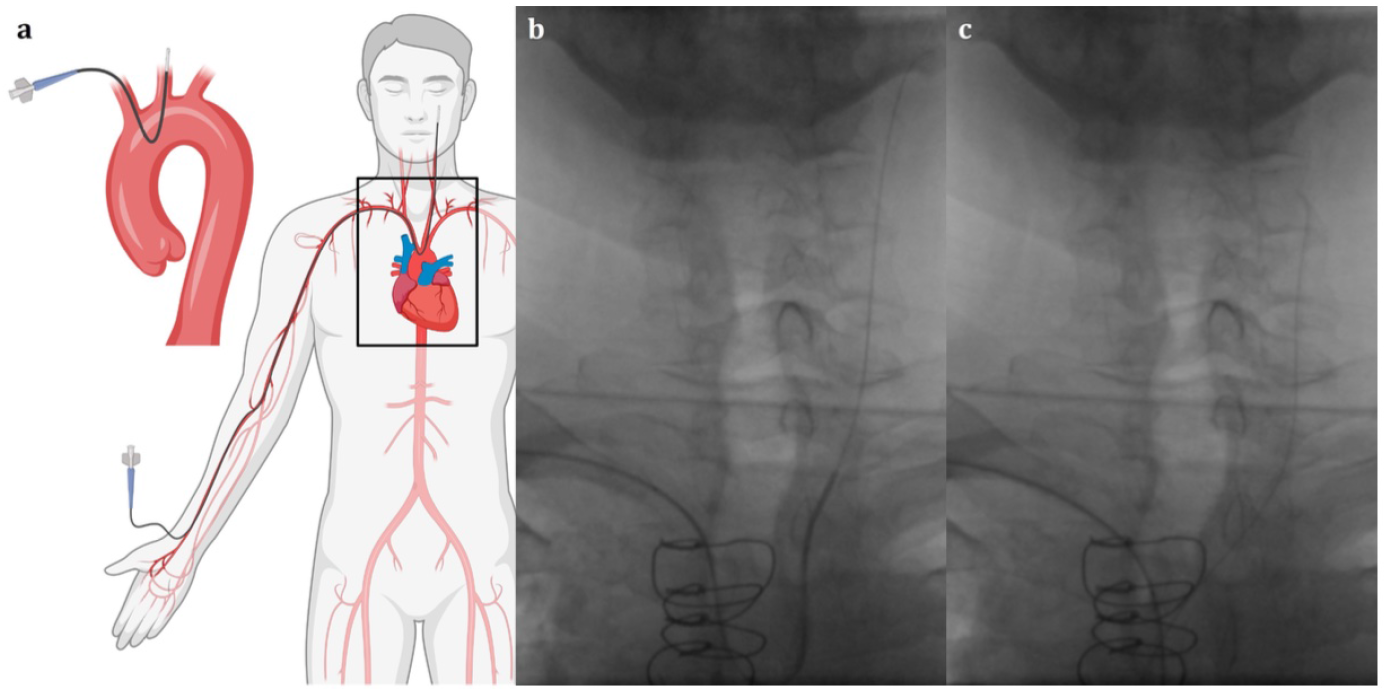
Herniation of a catheter-guidewire system. **(a)** Schematic depicting the tortuous anatomical path for transradial neurovascular access, highlighting the sharp bend through the aortic arch that the catheter-guidewire system must transit to access the brain (created with BioRender.com). **(b)** Image of a coaxial catheter-guidewire system prior to herniation. The guidewire is the thinner black line extending up the neck. The thicker black line extending from the left edge of the image is the catheter advancing coaxially over the guidewire. The top of the aortic arch is located near the bottom of the frame. **(c)** Image of the same catheter-guidewire system forming an extended loop and herniating down into the aortic arch after the surgeon attempted to advance the catheter along the guidewire up the left carotid. Stills taken from Supplementary Video 1.

Although interventionalists select a variety of devices based on personal preference and experience, no systematic mathematical framework for device selection currently exists [16–19], in part because the physics of herniation has not been established. We therefore explored the causes of herniation from an engineering perspective. We considered mechanical factors, which are well known to be critical for navigating complicated anatomy. Previous analysis of devices ranging from microcatheters to stents has focused on phenomenological engineering parameters including pushability, trackability, kink resistance, and durability [20, 21], and many international standards are in place for measuring the properties and efficacy of catheters [22–27] and guidewires [28]. However, based upon the role of bending evident from the above case study, we focused instead on flexural responses of the catheter-guidewire systems. Flexural rigidity, an indicator of resistance to bending, was a key consideration in even the first devices for endovascular surgery [29, 30]: low flexural rigidity is desirable for navigating complex arterial systems, but higher flexural rigidity is desirable for maintaining access during the introduction of additional devices. Catheters with a range of mechanical characteristics are available [31–34], including catheters with composite and spatially varying properties such as a soft distal tip [35–37]. A range of flexural rigidities for such devices has recently been characterized [38]. We hypothesized that the flexural rigidities of the catheter-guidewire system, combined with the patient-specfic anatomy of the pathway to be navigated, determine the likelihood of herniation.

Computational models already exist to predict how navigation and device advancement are associated with tissue deformation [39, 40], blood flow [41], and wall shear stresses [42], as well as to predict the effects of vascular surgery devices on hemodynamics [43]. Models also exist to estimate contact interactions between guidewires, catheters, sheaths, and support devices [44, 45]. These reveal that increased catheter diameter, impact angle, and interfacial area correlate with greater resistance during navigation [46]. However, the ways that the measurable properties combine with measurable features of patient specific anatomy to enable stable navigation and prevent herniation have not been explored. We therefore tested our hypothesis by developing a mathematical bifurcation model of catheter herniation, which we explored using computational and benchtop methods before applying it to cases of TRA surgery in human patients.

## 2 Results and Discussion

### 2.1 Catheter-guidewire navigation loss can be understood through a framework of bifurcation

To assess the stability of catheter-guidewire systems, we first developed a mathematical model of catheter navigation that treated the catheter-guidewire system as a thin, elastic line traversing the vasculature to a surgical site, and studied its potential bifurcation from this path (see Methods). Bifurcation, or rapid and uncontrolled change in configuration or path of the catheter-guidewire, is possible when the elastic bending energy stored in the catheter-guidewire system can be reduced by the system adopting a different shape or path within the vasculature [47].

To analyze the possibility of bifurcation during catheter navigation, we modeled the strain energy stored in a catheter-guidewire system traversing the aortic arch in a TRA procedure. In this procedure, the catheter-guidewire system follows the radial artery from the wrist and up the arm, traversing the inominate before making a bend with a substantially tighter radius of curvature to pass through the aortic arch and up into the left carotid that leads to the brain (Fig. 1). If the navigation is successful, the catheter-guidewire system has a radius of curvature of 2*R* as it traverses the aortic arch (Fig. 2a). The mechanical energy stored per unit length in that region scales as scales as the square of the curvature, *κ*^2^ = (1*/R*)^2^ [47].

**Fig. 2.**
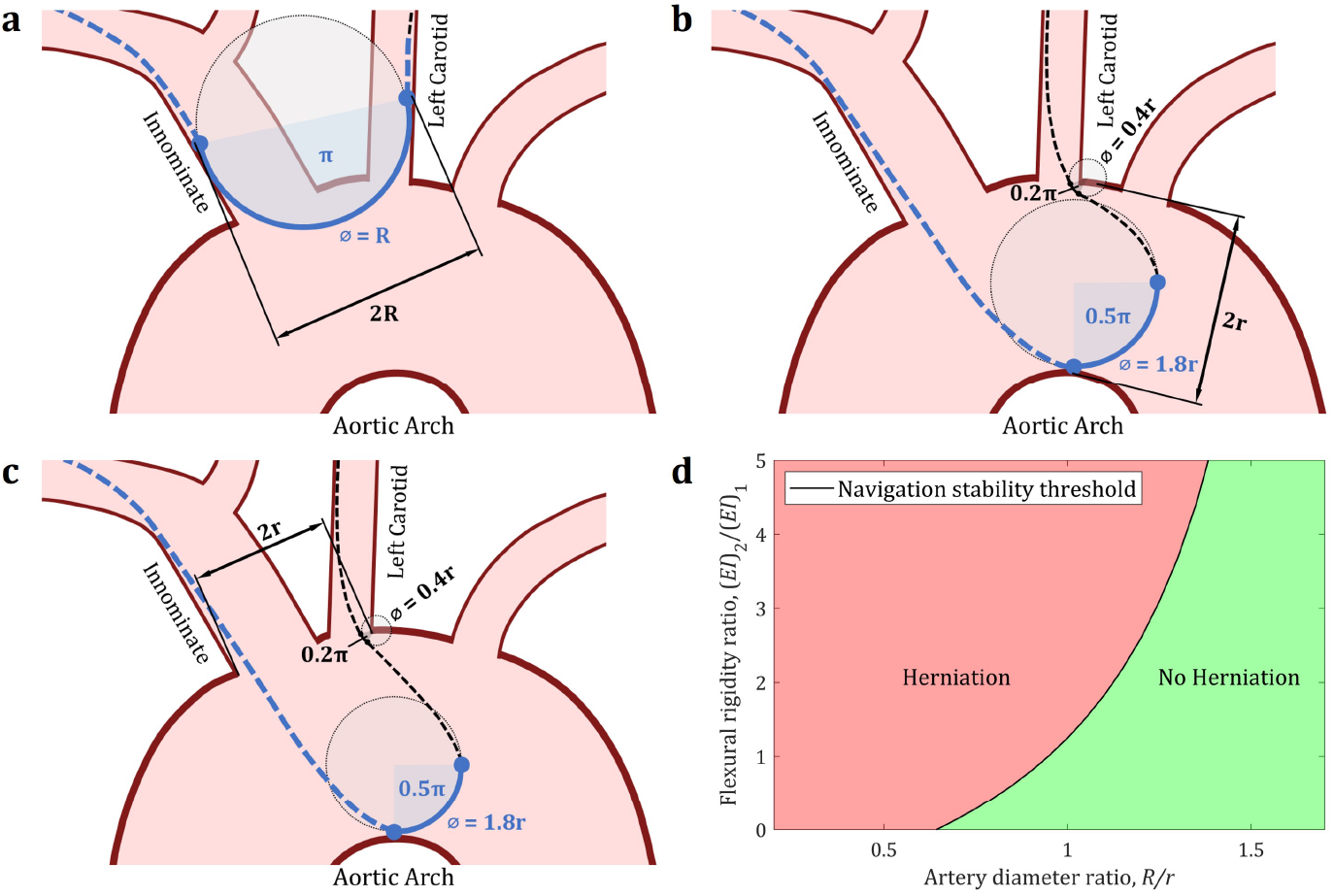
A mathematical model of endovascular navigation stability. **(a-c)** Schematics of successful (a) and unsuccessful (b-c) navigation of a catheter-guidewire system. Black lines represent the first device. Blue lines represent the coaxial combination of the first and second devices. Solid lines represent regions of high curvature included in the energy calculations, while dashed line represent regions with minimal curvature that are neglected in the energy calculations.**(a)** Schematic of a successful transit of the aortic arch. **(b)** Schematic of a herniated state in the case where the critical dimension, 2*r*, is given by the aortic arch diameter. **(c)** Schematic of a herniated state in the case where the the aortic arch is relatively wide compared to the spacing between the innominate and left carotid, so that the critical dimension, 2*r*, is instead given by the distance between the base of the innominate artery and the left carotid artery. **(d)** A phase diagram defined by the theoretical navigation stability criterion showing the ratios of device flexural rigidities that result in herniation for specific patient anatomies.

If herniation occurs, the catheter-guidewire system drops from the carotid artery, forming a tighter loop within the aortic arch (Fig. 2b,c). Several possibilities exist for the path of this catheter-guidewire system trajectory, each of which necessarily requires tighter bends of the catheter-guidewire system within the aortic arch, resulting in a higher strain energy per unit length. However, this higher strain energy per unit length can be stored within a region of shorter length in the catheter-guidewire system, potentially resulting in a net reduction of energy. Following the principle of minimum potential energy [47], any assumed shape that can exist in static equilibrium will provide an upper bound on the energy stored by the catheter-guidewire system in the herniated state. Bifurcation is possible if the energy of any of these assumed, herniated catheter-guidewire system shapes is lower than that associated with successful navigation.

When multiple devices are passed coaxially along acute bends in the vasculature, following this approach leads to a mathematical navigation stability criterion that predicts conditions under which herniation is not predicted to occur:

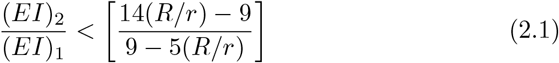

where (*EI*)_1_ and (*EI*)_2_, respectively, represent the flexural rigidities of the first device (typically a guidewire) and the second device that is passed coaxially over or through the first. The values of (*EI*)_1_ and (*EI*)_2_ can be selected by the surgeon, and the value of *R/r* is patient-specific. A second device with sufficiently small flexural rigidity (*EI*_2_) for the inequality in Eq. 2.1 to hold is predicted to successfully navigate the bend, while a flexural rigidity ratio greater than the quantity on the right side of that equaation is predicted to herniate. For a neurointerventional surgery using TRA, the quantity 2*R* is the distance between the outer walls of the innominate and the left carotid over which the loop with the lowest curvature (1*/R*) will form if the system successfully transits the aortic arch (Fig. 2 a). The quantity 2*r* is the limiting distance over which looping can occur in the assumed herniated state, such as the diameter of the aortic arch (Fig. 2b) or, if the aortic arch is relatively wide and the innominate and left carotid are relatively close together, the distance between the base of the innominate artery and the left carotid artery (Fig. 2c). The navigation stability criterion defines a curve in a phase diagram (Fig. 2d) that predicts the patient-specific combinations of tools for which successful navigation is likely. The two state variables are the flexural rigidity ratio, (*EI*)_2_*/*(*EI*)_1_, between the two coaxial devices, and the patient-specific artery diameter ratio, *R/r*. Published values of (*EI*) are available for a range of surgical devices [38]. A specific patient will have, based upon vascular anatomy, a single value of the artery diameter ratio. If the artery diameter ratio is below a threshold of approximately 0.6, successful navigation is not possible with any combination of devices and the surgeon must choose a different approach to the brain. For patients with greater artery diameter ratios, the surgeon may choose catheter-guidewire systems with increasingly high flexural rigidity ratios.

### 2.2 Computational verification of the navigation stability criterion

We first tested the basic assumptions underlying the analytical navigation stability criterion by comparison to a computational (finite element) model of catheter navigation. Two key assumptions were tested. We asked whether the assumed shapes used to derive the navigation stability criterion were adequate to predict herniation of an unconstrained catheter-guidewire system traversing an idealized TRA surgical path (Fig. 3). We also asked whether the navigation stability threshold, which evaluates a catheter and guidewire together, would serve as a valid bound for the actual clinical procedure, involving a guidewire that has already navigated the surgical path and a catheter that is being advanced gradually over the guidewire.

**Fig. 3.**
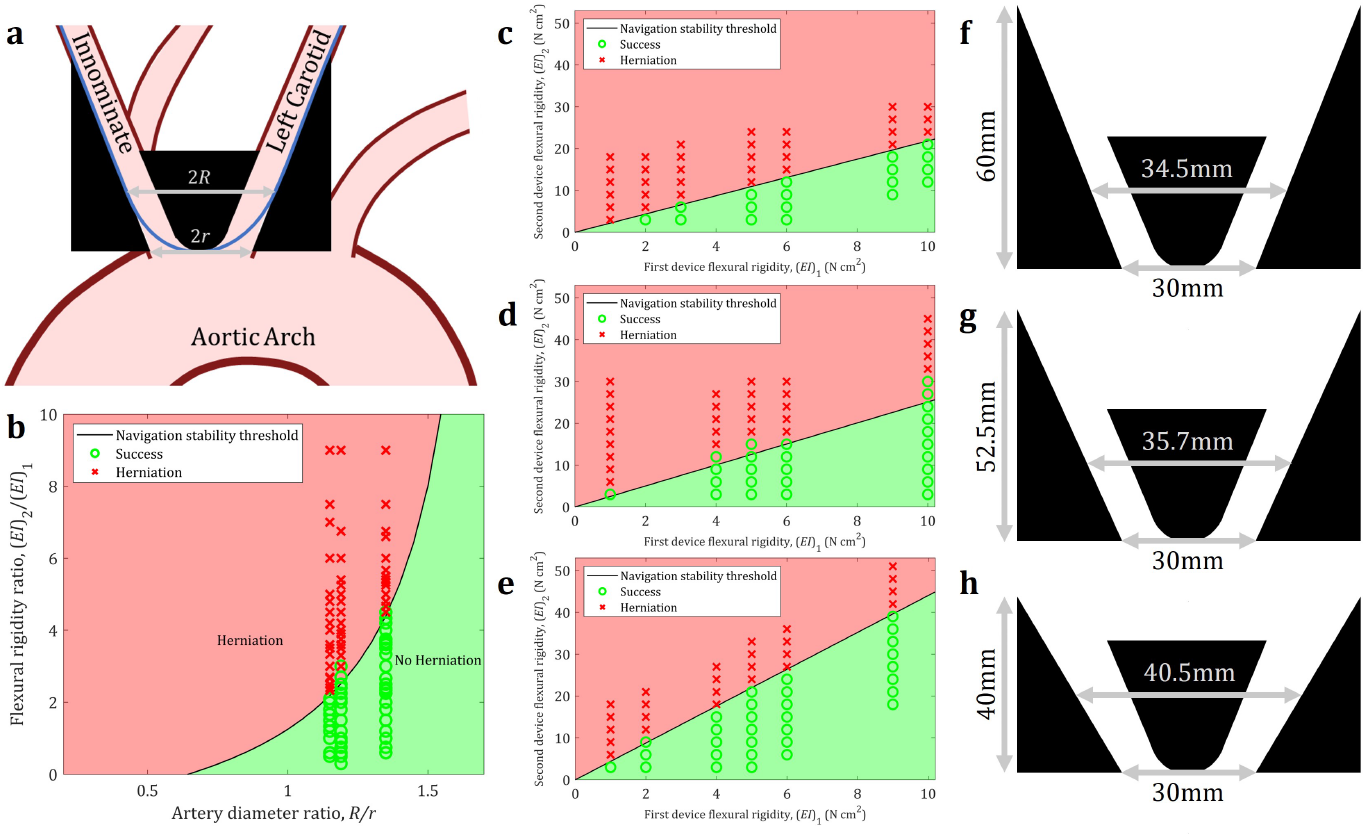
Computational verification of model assumptions. **(a)** Schematic showing the spatial domain of a finite element model (in black) superimposed on the aortic arch. The dimension 2*R* is the diameter of the largest loop that a coaxial catheter-guidewire system (depicted in blue) can form while successfully transiting the arch. This diameter was measured based from a simulation of a successful transit. The dimension 2*r* is the distance between the base of the rigid bodies representing the outer walls of the innominate and the left carotid arteries (the shortest, limiting distance over which herniation can occur in this case). **(b)** The simulated herniation outcomes of 160 simulations of unique combinations of device flexural rigidity ratio and arch geometry overlaid on the navigation stability diagram. **(c**,**d**,**e)** The same data, instead plotted by first and second device flexural rigidity overlaid on separate phase diagrams for each of the three computational models. The slope of the phase boundary decreases for narrower arch geometry. **(f**,**g**,**h)** Schematics showing the critical dimensions of each of the three computational models studied. Each model is associated with the adjacent phase diagram. See Supplementary Data 1 for raw data.

A two-dimensional (2D) model of idealized, rigid, frictionless vasculature and a relatively large aortic arch was considered. By varying the device flexural rigidities as well as the geometric dimensions of the arch, we simulated 160 unique TRA procedures to determine whether bifurcation and hence herniation occurs in each specific case. Equations were solved using elements that modeled Euler-Bernoulli beams and rigid boundaries in the Abaqus finite element package (Dassault Syst‘emes Simulia Corp., Vélizy-Villacoublay, France; see Methods).

Results demonstrated that herniation of a catheter-guidewire system is a bifurcation event (Fig. 3b). In each simulation, the guidewire (first device, flexural rigidity (*EI*)_1_) was initially placed in an equilibrium state over the surgical pathway, and a catheter (second device, flexural rigidity (*EI*)_2_) was tracked over the first device, advanced from the left arterial branch (representing the innominate) through the arch to the second arterial branch (left carotid). For each of the three patient anatomies considered, represented by three different artery diameter ratios, bifurcation occurred for higher values of the flexural rigidity ratio, but not for lower values (Fig. 3a). Bifurcation was identified as a deviation from the surgical path that prevented the catheter from navigating successfully up the left carotid while simultaneously drawing the guidewire down from its initial position in the carotid. Some simulations became unstable numerically during this process.

For a given patient anatomy, as represented by the artery diameter ratio, the flexural rigidity ratio of the catheter-guidewire system determined navigational success. As predicted by the model, tighter bends (smaller *R*) in the anatomy led to herniation at lower values of the flexural rigidity ratio (Fig. 3b). As expected for a model that serves as an upper bound, the navigation stability threshold predicted trends correctly, and was slightly conservative: a few cases of successful navigation occurred in the computational simulations for conditions that were slightly beyond those predicted to cause herniation, but no cases of unsuccessful navigation occurred for conditions predicted to not cause herniation Fig. 3c,d,e). Overall, the navigation stability criterion was predictive of herniation in the numerical model (Fig. 3b), with a sensitivity of 100.0%, specificity of 91.9%, positive predictive value (PPV) of 93.5%, and negative predictive value (NPV) of 100.0%. Results revealed that the navigation stability criterion could accurately predict the critical threshold in a simplified, 2D problem that accounted for the intermediate stages of advancing a catheter over a guidewire.

### 2.3 *Ex vivo* verification of the navigation stability criterion

The 2D analyses are expected to serve as upper bounds to the 3D problem, because energy stored in herniated catheter-guidewire systems can be further reduced by out-of-plane motion. The analyses from above also neglect friction and fluid within the vasculature because they do not affect the relationship between the shape of the catheter-guidewire system and the energy stored in it. To verify these assumptions, we constructed a rigid, 3D printed vascular model of the aortic arch based on a representative computed tomography (CT) scan of anonymized patient data. The 3D printed model was immersed in tap water, and a total of 117 unique combinations of commercially available catheters and guidewires were then advanced through the 3D printed model.

The resulting plot of herniation outcomes again demonstrates the expected bifurcation behavior (Fig. 4c). The likelihood of herniation again increased with increasing flexural rigidity of the second device. The threshold for herniation increased with increasing flexural rigidity of the first device. In terms of predictiveness *ex vivo*, applying the navigation stability criterion yielded a sensitivity of 92.3%, specificity of 86.5%, PPV of 89.6%, and NPV of 90.0%. Results demonstrated that the navigation stability criterion was predictive of herniation even in a 3D model with fluid and friction, and verified the assumption that out of plane twisting and frictional effects have only minor influence on the likelihood of herniation.

**Fig. 4.**
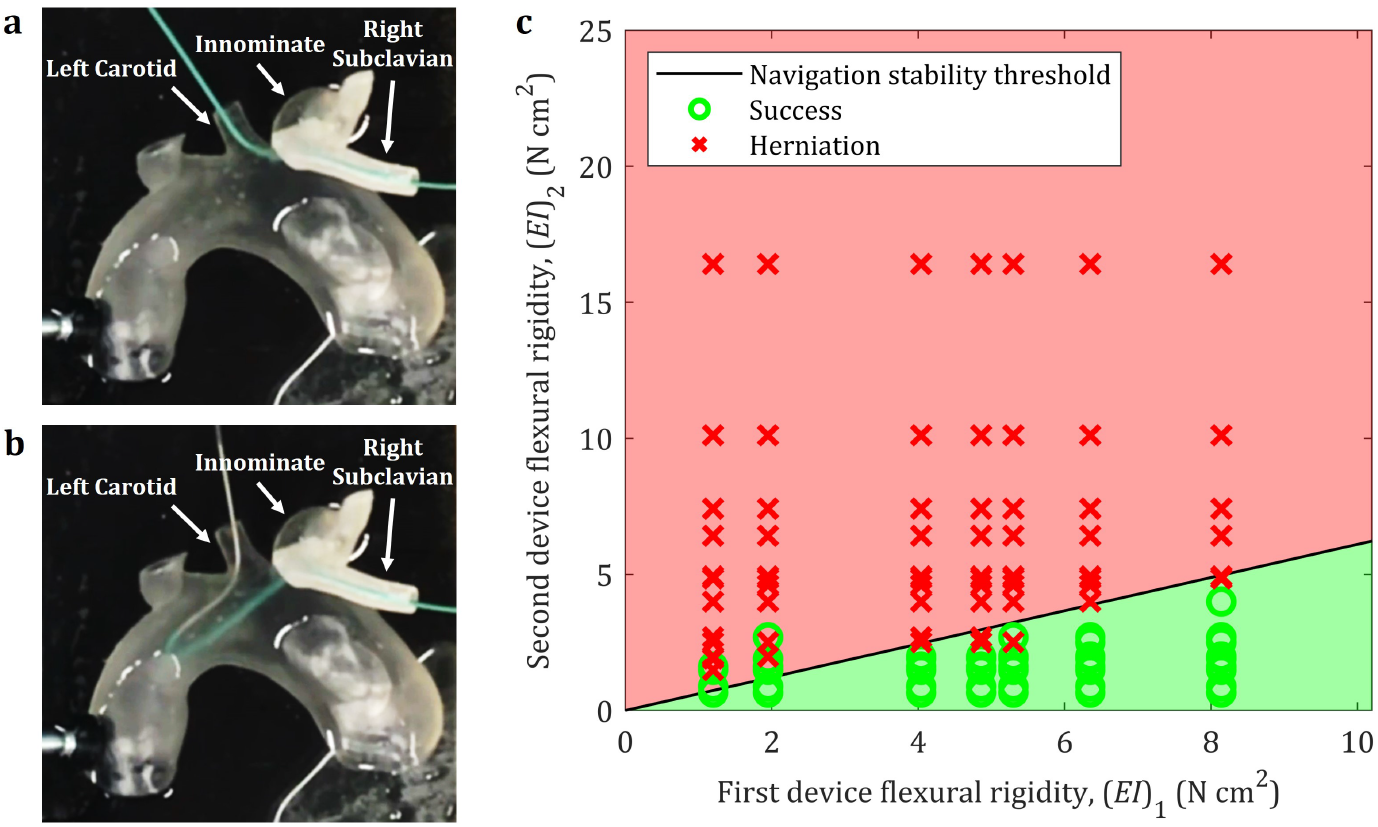
Benchtop verification of the navigation stability criterion. Representative photos of **(a)** successful transit and **(b)** herniation recapitulated in a 3D printed, patientspecific, rigid, benchtop *ex vivo* model. **(c)** The herniation outcomes of the 117 unique coaxial device combinations tested in the benchtop model overlaid on the phase diagram specified by the navigation stability criterion for the aortic arch dimensions of the 3D printed model. See Supplementary Data 2 for raw data from these experiments.

### 2.4 *In vivo* verification of the navigation stability criterion

We evaluated the clinical applicability of the navigation stability criterion by retrospectively applying it to previous neurovascular surgical cases performed in the Washington University Department of Neurosurgery. A total of 11 cases were evaluated, 6 of which were completed successfully, and 5 of which involved herniation. For each patient, the artery diameter ratio was measured from preoperative CT angiography (CTA) images, and the surgical device pairings and surgical outcome (success versus herniation) were determined based upon the procedure notes.

In each case, the first device or device pair was advanced via a TRA approach, and navigated through the aortic arch from the innominate to the left carotid (Fig. 5). Plotting these results on top of the phase diagram specified by the navigation stability criterion demonstrates that the surgical outcomes followed the expected bifurcation behavior (Fig. 4e). In the case of the patient data, the navigation stability criterion was again highly predictive of herniation *in vivo*, with a sensitivity of 100.0%, specificity of 85.7%, PPV of 80.0%, and NPV of 100.0%. The navigation stability criterion remained predictive of herniation in spite of all of the complexities associated with actual surgery (variable blood flow, flexible vessel walls, vessel wall movement, etc.), suggesting its usefulness in the clinical setting.

**Fig. 5.**
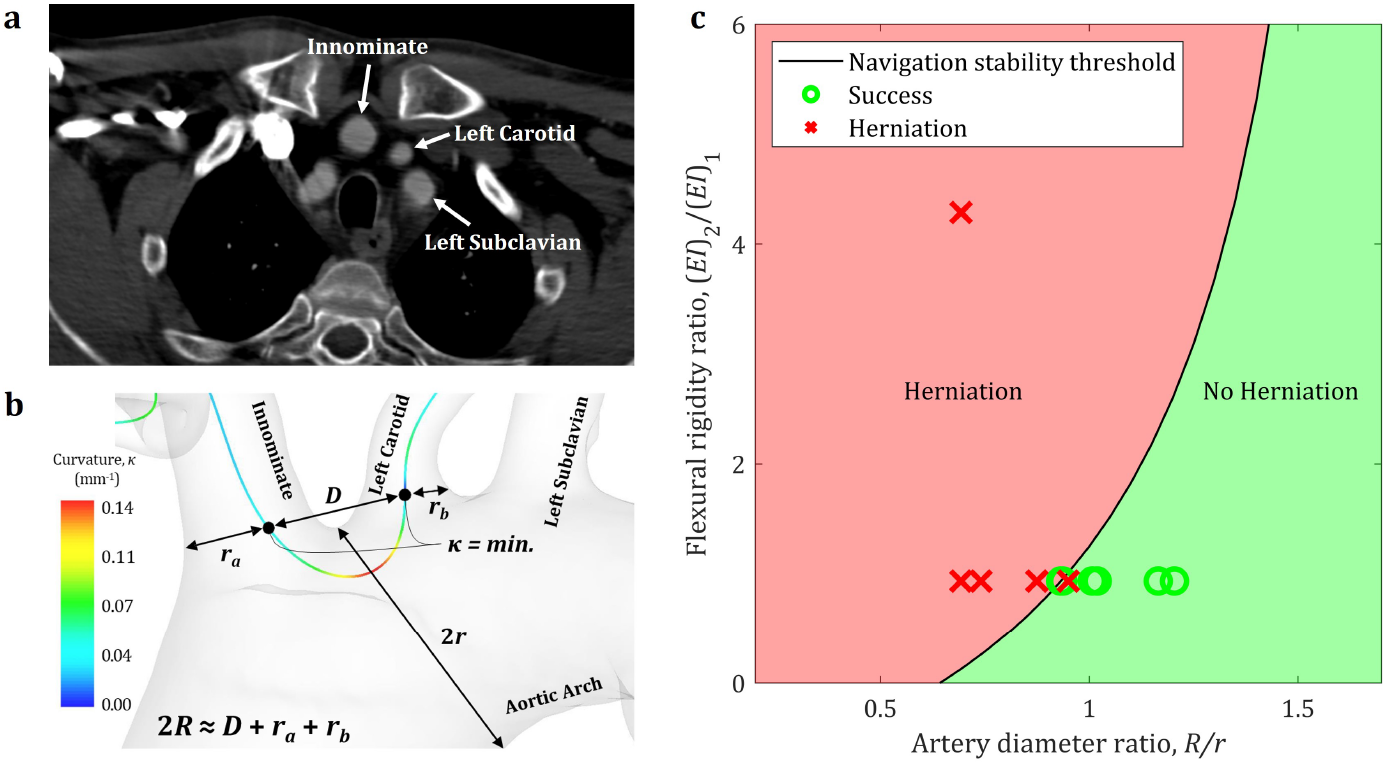
*In vivo* application of the navigation stability criterion. **(a)** Cross-section of a representative patient CT angiogram captured prior to surgery. The slice shown is at the level where the innominate, left carotid, and left subclavian arteries intersect the aortic arch (as labeled) **(b)** Schematic showing the key patient-specific arch dimensions measured from a representative three-dimensional surface model generated by segmenting a patient CT angiogram. **(c)** Navigation outcomes from 11 surgical cases plotted based on the measured, patient-specific arch dimensions and flexural rigidities of the devices used, overlaid on the phase diagram specified by the navigation stability criterion. See Supplementary Data 3 for the raw data used in the retrospective patient analysis

### 2.5 The navigation stability criterion shows potential application to preventing herniation in surgical procedures

We verified the navigation stability criterion across multiple experiments with increasing fidelity. While overall accuracy decreased slightly with increasing fidelity, the criterion was still highly predictive in all cases. In combination, the results demonstrated that the criterion is robust across conditions and a range of possible vascular geometries. The current work underscores the key contribution that surgical tool selection plays in preventing endovascular catheter herniation. Currently, surgeons rely on qualitative experience and intuition to determine which combination of devices to use. The navigation stability criterion developed here provides a reliable, quantitative metric which surgeons can use to inform their tool selection for a specific patient anatomy. With two simple measurements of the patient’s anatomy made ahead of surgery, the navigation stability criterion may allow surgeons to avoid herniation, reducing procedure time and potential complications. Because the needed preoperative computed tomography scans are typically performed for all patients undergoing endovascular surgical procedures, no additional procedures have to be added to the current workflow to apply the navigation stability criterion.

Only a few devices are currently used for TRA, and even fewer are marketed specifically for TRA, as evidenced by the limited number of device combinations present in our sample of surgical cases (Fig. 4e). The navigation stability criterion can possibly guide development of catheter-guidewire systems that successfully transit specific patient anatomies not only for transradial, neurovascular surgeries but also for endovascular procedures more broadly.

We note that the study has several limitations. The method used to assess device flexural rigidity quantifies only the stiffest portion of the device and fails to capture variations in stiffness along its length. This may be an issue for catheters and guidewires with significant transition zones. Also, the navigation stability criterion derived here will only apply to standard Type I, II, and III aortic arches, where the innominate artery and the left common carotid artery have distinct origins on the aortic arch. Other arches, such as arteria lusoria and bovine arch types in which the left common carotid branches off the innominate artery, have significantly different vessel boundaries and thus involve distinct herniation states, requiring different definitions of the successful transit and heriated states. Finally, the navigation stability criterion derived here only applies to bi-axial systems (those involving two devices). However, extending to three (tri-axial) or more devices simply requires adding additional energy terms.

## 3 Conclusions

Our results show that endovascular catheter herniation is a bifurcation phenomenon that occurs when a catheter-guidewire system forms an extended loop that prevents the successful transit of the catheter when an interventionalist attempts to navigate it around a tight bend. The navigation stability criterion derived here provides a quantitative metric predicting the likelihood of herniation based on measurable, patient-specific vascular geometry and device mechanical properties, and was accurate for a range of model tests and surgical cases. Interventionalists currently rely on experience, intuition, and catheter preference to avoid herniation. These results instead offer a physics-based framework for identifying the threshold for catheter herniation, and provide a quantitative and objective basis for device selection during endovascular procedures.

## 4 Methods

### 4.1 Navigation stability criterion

To quantify a navigation stability criterion, we considered the elastic bending energy associated with the successful transit of the arch and the herniated state. Assuming that the catheter-guidewire system tends to adopt the state with the lowest internal energy, we derived a navigation stability criterion that predicts the critical threshold based on the patient’s vascular geometry and the flexural rigidities of the devices used.

The model neglected friction. Most devices used in endovascular surgery are coated with lubricious films designed to minimize friction [48]. This results in frictional energy dissipation on the order of 0.1 J/m [48, 49], while elastic energy storage due to bending is on the order of 1 J/m [49]. Since the energy associated with bending dominates, we limited our analysis to consider only the elastic bending energy of the devices.

To calculate these energies, we first defined each state based on the positions of the vessel boundaries and by analyzing videos of herniation (Fig. 1 and Supplementary Video 1). For the transradial case, device bending occurs mostly in a single plane. Thus, to simplify the problem, we limited the analysis to two-dimensions.

In the successful transit state, the catheter-guidewire system loops down from the innominate artery, through the aortic arch, and up the carotid artery, forming the largest curvature loop possible in order to minimize bending energy (Fig. 2a). In the herniated state, the devices form an extended loop, with one bend in the aortic arch followed by a second, smaller bend around the corner where the carotid intersects the arch (Fig. 2b,c). We assumed that the energy stored in the regions connecting these two bends possess negligible curvature, and thus neglected their energy contributions. We also assume that the system is stationary in both the successful transit and herniated states. Note that these definitions likely give an upper bound for the energy in both states; the actual energies would be expected to be lower due to minor, energetically favorable reconfigurations of the catheter-guidewire system.

Modeling both devices as a beam, the internal energy stored due to bending is given generally by the Euler-Bernoulli beam theory as follows:

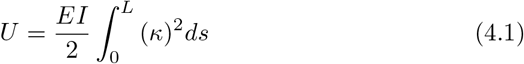

where *U* is internal stored elastic energy, *E* is the elastic modulus, *I* is the second area moment of the area of the cross section, *κ* is the neutral bend curvature, and *L* is the length of the bend. The combined term *EI* is the flexural rigidity, which quantifies a beam’s resistance to bending.

Applying Eq. 4.1, the the energy *U*_*s*_ associated with the successful transit state can be calculated as follows:

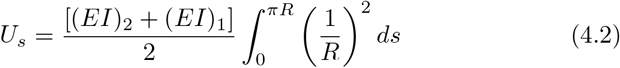

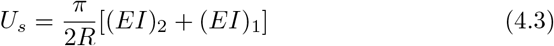

where *EI* and *k* are as previously defined, *R* is as defined in Fig. 2a, and devices 1 and 2 are the first and second devices used coaxially in the system, respectively.

Similarly, the energy *U*_*h*_ associated with the herniated state is given by:

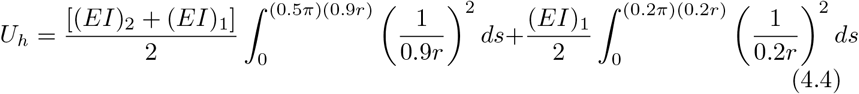

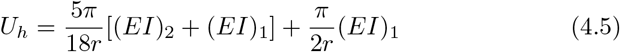

where *EI* and *k* are as previously defined, *r* is as defined in Fig. 2b,c, and devices 1 and 2 are the first and second devices used coaxially in the system, respectively.

To avoid herniation, the energy associated with the successful transit of the arch must be lower than that of the herniated state:

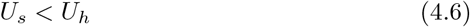

Substituting in the expressions for *U*_*s*_ and *U*_*h*_:

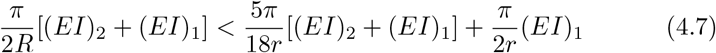

Simplifying gives the final navigation stability criterion:

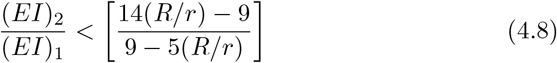

### 4.2 Computational herniation analyses

A quantitative finite element model was developed in the Abaqus environment to replicate the passage of a second device coaxially along a placed device through a tortuous bend in the aortic arch. We simulated the bend connecting the right innominate artery to the left carotid artery in two dimensions to mimic right TRA for neurointerventional access. In order to simplify the computational analysis, we chose to exclude the bottom of the aortic arch from the Abaqus model. This effectively reduces the number of contacts that must be simulated and restricts the possible herniation states to that depicted in Fig. 2c.

Three rigid body parts were drawn and positioned to create a bend connecting the two arteries. Coarse patterns of quadrilateral-dominated elements were meshed onto each region (65-88 elements per body). Three iterations of the arch design were modeled with varying vessel angle by changing the height of the left and right artery boundaries from 60 mm to 52.5 mm and 45 mm (Fig. 3f,g,h). The general dimensions and scale of the arch design were modeled in accordance with a standard Type I arch (Fig. 3a).

The device bodies were modeled in Abaqus using wire segments meshed with 80 beam elements. A 40 cm wire was constructed and split into two colinear sections: a 25 cm flexible section and a 15 cm stiff section. The elastic modulus of each section was modulated throughout the study to change the desired flexural rigidity. The beam segment was built with a 2 mm outer diameter tubular profile with 0.3 mm thickness to approximate a 6 Fr device.

The arch boundaries remained rigid throughout the study, and the device was steered by displacing the proximal tip of the system and leaving the distal shaft free. Contact interactions were imposed between the artery walls and the sides of the device. The middle arch body was designed to interact with the top of the devices, and the left and right outer artery walls with the bottom of the devices. In the initial state, the two device system was positioned on the cranial side of the simulated left carotid artery. The entire catheter system was then pulled through the arch by incrementally shifting the proximal point. Once in place, the elastic modulus was adjusted for each section to match the desired flexural rigidity for the device pairing. The proximal end of the second device was displaced linearly towards the arch until the its distal tip fully transited the arch.

Since herniation is a bifurcation event characterized by large qualitative translations, the deformed result of herniation is challenging to quantify, especially using numerical methods. We assessed herniation based on two output parameters: the simulation output (pass or abort) and the deformed result (success or herniation). The computational simulation indicated when the simulation of catheter passage became unstable, as arises during bifurcation, and aborted the analysis. A qualitative assessment was also performed to determine if the second device had fully transited the arch in the deformed final state (see Supplementary Figure 1). This testing process was repeated 160 times, varying the device and geometric combinations; 50, 54, and 56 simulations were performed on the 60 mm, 52.5 mm, and 45 mm tall arch models, respectively.

For assessment of the model, the ratio of *R/r* was measured from the simulation output using ImageJ. Representative images were collected for each arch type of successful passages that passed the full study. The dimension 2*R* was measured between the highest nodes maintaining the curvature of the bend that do not touch the artery wall, and 2*r* was measured between the two lowest central nodes of the left and right rigid body artery walls (Fig. 3a).

### 4.3 Patient data collection and computed tomography scan protocol

Patient data were collected restrospectively from neruovascular surgical procedures conducted by the Washington University School of Medicine Department of Neurosurgery. The study protocol was approved by the Washington University Institutional Review Board (IRB), and informed consent was waived given the retrospective nature of the study and that no identifying information is required for analysis.

Pre-procedure head and neck CTA images were analyzed from 11 adult patients undergoing diagnostic angiography using transradial access. Patients with standard arch types 1, 2, 3, and bovine arch variants with a common origin for the innominate and left common carotid arteries were included. Patients with arteria lusoria or bovine arch variants in which the left common carotid artery originates separately from the innominate artery were excluded.

Contrast medium with an iodine content of *>*300 mg/mL (Optiray 350; Mallinckrodt Pharmaceuticals, Staines-upon-Thames, United Kingdom) was administered at 37^*°*^C through a 20G peripheral arterial catheter in the radial or femoral artery at a flow rate of 4-5 mL/s. CTA was performed using either a 16-slice or a 64-slice multidetector scanner. Serial CTA imaging at the level of the pulmonary artery and the ascending and the descending aorta were obtained at 1.0-s intervals from 0 to 99 s. The following scan parameters were applied for the 16-slice and 64-slice scanners, respectively: collimation 0.75/0.63 mm, pitch 0.25/0.24 mm, reconstruction slice thickness 1.0/0.63 mm, increment 1.0/0.63 mm.

The surgical devices used and the success vs. herniation outcome for each of the 11 patient cases were determined based on the note for the procedure.

### 4.4 Design of rigid, 3D printed aortic arch model

To create a 3D printed model of the aortic arch, the CTA images from a patient with a standard Type I arch were selected. CTA images with a slice thickness of 1.0 mm were segmented and visualized in a commercial processing and editing software (3D Slicer, www.slicer.org). A volume of interest ranging axially from above the brachial artery to below the middle cerebral artery (MCA) was resliced to generate iso-cubic voxels of dimensions 0.500 *×* 0.500 *×* 0.500 mm^3^. The aortic arch and major branches (innominate artery, left common carotid artery, and left subclavian artery) were segmented using a semi-automatic threshold-based region growing command. The segmented geometry was then edited to remove branches that were poorly visualized or not under investigation. A three-dimensional (3D) surface mesh was generated and exported as a stereolithography (STL) file. STL processing was performed on Mesh Mixer (Mesh Mixer 2.8; Autodesk Inc., San Rafael, CA, USA) to smooth the surface and correct errors in the mesh. A wall thickness of 1.5 mm was digitally produced, and the space occupied by the lumen was subtracted to create primary hollow models for stereolithography apparatus (SLA) 3D printing.

The model was printed using a transparent resin with the consistency of hard plastic. Upon removal of support structures, the SLA-printed model was smoothed using grade #0000 steel wool (Rhodes American Steel Wool, Super Fine Grade #0000 pads; Homax Products, Inc., Bellingham, WA, USA) followed by wet sanding with 3000 grit sandpaper (3M Wetordry Sanding Sheets; 3M, Saint Paul, MN, USA). The powder residue was removed with a standard multi-surface cleaner and disinfectant spray and subsequently rinsed in water. The critical dimensions *R* and *r* were then measured manually off of the 3D printed model, accounting for wall thickness (see Supplementary Figure 2 for model dimensions).

### 4.5 Benchtop herniation experiments

Catheter herniation was recapitulated in benchtop experiments using the rigid, 3D printed vascular model. Briefly, catheters with outer diameters of 1.667 to 2.000 mm were advanced across a single bend from the innominate artery branch through the aortic arch and out the left common carotid artery branch, so as to simulate access to the left anterior circulation of the brain vasculature from the right radial artery. Guidewires with outer diameters of 0.635 to 0.889 mm were then advanced coaxially through the pre-positioned catheter, and the success vs. herniation outcome was noted (see Supplementary Table 2 for full list of devices and Supplementary Video 2 for a demonstration of the procedure).

### 4.6 Geometric analysis of patient vasculature

All image processing, surface mesh generation, and geometric analyses were performed using the Vascular Modeling Toolkit (VMTK) library. The aortic arch, innominate artery, left common carotid artery, and left subclavian artery were manually segmented using the level sets method. The zero set level surface was extracted using the marching cubes algorithm and smoothed using a non-shrinking Taubin filter. A loop subdivision scheme was used to generate a higher order surface model. Centerlines were computed from the innominate artery at its proximal end up to the common carotid artery at its distal end, and from the ascending aortic arch up to the descending aortic arch. Centerlines were sampled at intervals of 0.25 mm with a smoothing factor of 0.7 and 500 iterations. Point-wise curvature, torsion, maximal inscribed sphere radii, and corresponding coordinates in three dimensions were calculated along centerlines, as provided by the VMTK library, and extracted for further analysis in MATLAB (The Mathworks, Natick, MA). Geometric parameters were obtained using a standardized protocol for TRA vascular dimension measurements as depicted in Fig. 4d.

### 4.7 Flexural rigidity assessment of guidewires and catheters

Until recently, vascular surgeons had no quantitative metric by which to assess the mechanical, bending performance of the surgical devices at their disposal. Hartquist et al. provided a framework for quantifying the flexural rigidity of catheters and catalogued values for common commercial devices [38]. The methods from this work were replicated to evaluate the flexural rigidity of the guidewires and catheters used in the analyses above (see Supplementary Tables 2 and 3, respectively, for the flexural rigidities of the devices used in the benchtop study and patient cases).

## Supplementary information

**Supplementary Video 1**

Representative video of catheter herniation *in vivo*.

**Supplementary Video 2**

Representative video of catheter herniation *ex vivo*.

**Supplementary Figure 1**

Four representative outputs from the computational simulations.

**Supplementary Figure 2**

Dimensioned rendering of the arch selected for the 3D printed, benchtop model.

**Supplementary Table 1**

Raw data from the computational analyses.

**Supplementary Table 2**

Raw data from the benchtop navigation experiments.

**Supplementary Table 3**

Raw data used in the retrospective patient analyses.

**Supplementary Table 4**

Catheter, sheath, microcatheter, and guidewire flexural rigidity measurements for common commercial devices.

## Declarations

### Funding

This work was supported by Society for Vascular Surgery Foundation Research Investigator Award (M. Zayed), American Surgical Association Research Fellowship Award (M. Zayed), K08HL132060 (M. Zayed), Center for Innovation in Neuroscience and Technology (E.C. Leuthardt), the National Science Foundation Science and Technology Center for Engineering MechanoBiology (grant CMMI 1548571; G.M. Genin), and the National Institutes of Health (R41HL150963, M. Zayed, and G.M. Genin).

### Competing interests

M.A.Z. and G.M.G. have financial interest in Caeli Vascular, Inc. M.A.Z., J.W.O., and G.M.G. have financial interest in Inflexion Vascular, LLC. J.W.O. provides consulting services for Medtronic, Microvention, Terumo, and InNeuroCo.

### Ethics approval

Study protocol was approved by the Washington University IRB.

### Consent to participate

Not applicable.

### Consent for publication

The views expressed in this article do not necessarily reflect those of The Aerospace Corporation.

### Availability of data and materials

The datasets generated during and/or analyzed during the current study are available from the corresponding author upon reasonable request.

### Code availability

Author-generated code are available from the corresponding author upon reasonable request.

### Authors’ contributions

C.M.H., J.V.L., V.C., H.R.L, M.A.Z., J.W.O., and G.M.G. conceived the study design. C.M.H., J.V.L., M.Y.Q., and C.S. performed data analysis and interpretation of the data. C.M.H., J.V.L., and M.Y.Q. drafted the manuscript. M.A.Z., J.W.O., and G.M.G. supervised the project. All authors revised and approved the final manuscript.

