## supplementary figures for "Stability of navigation in catheter-based endovascular procedures"

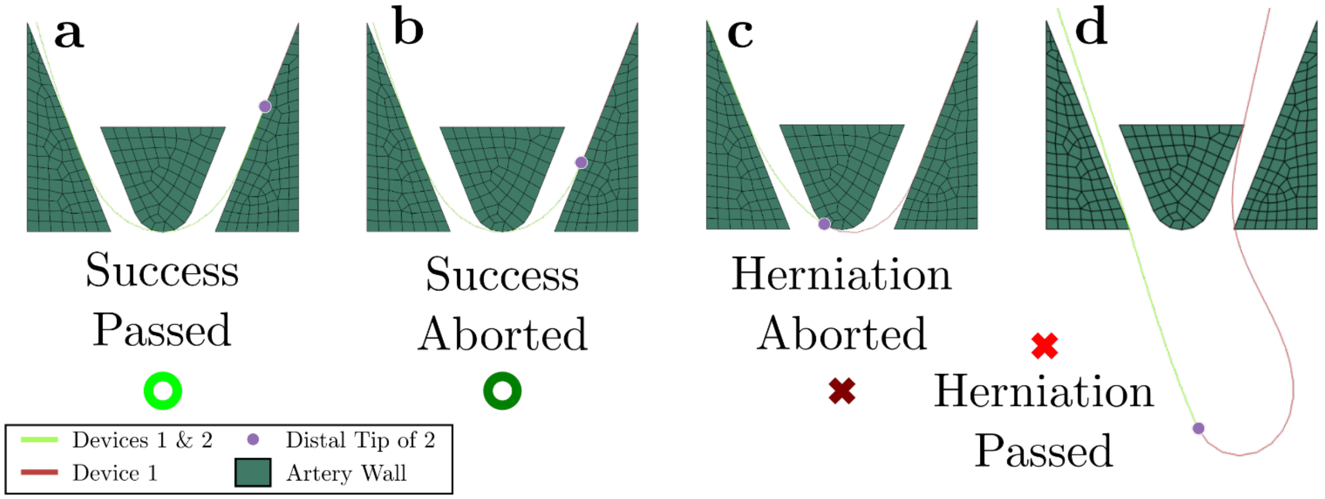


**Supplementary Figure 1.** The four representative outputs from the finite element herniation study in Abaqus. Success vs. herniation was evaluated qualitatively based on the simulation output. **(a)** Devices 1 and 2 both successfully transit the bend, and the simulation runs to completion. **(b)** The devices successfully transit the bend, but the simulation aborts before completion. Occurred in 12% of success cases. **(c)** The simulation aborts before device 2 transits the bend. This output was considered a herniation and made up 89.5% of herniation cases. **(d)** The simulation runs to completion, but the second device fails to transit the bend. Occurred in 10.5% of herniation cases

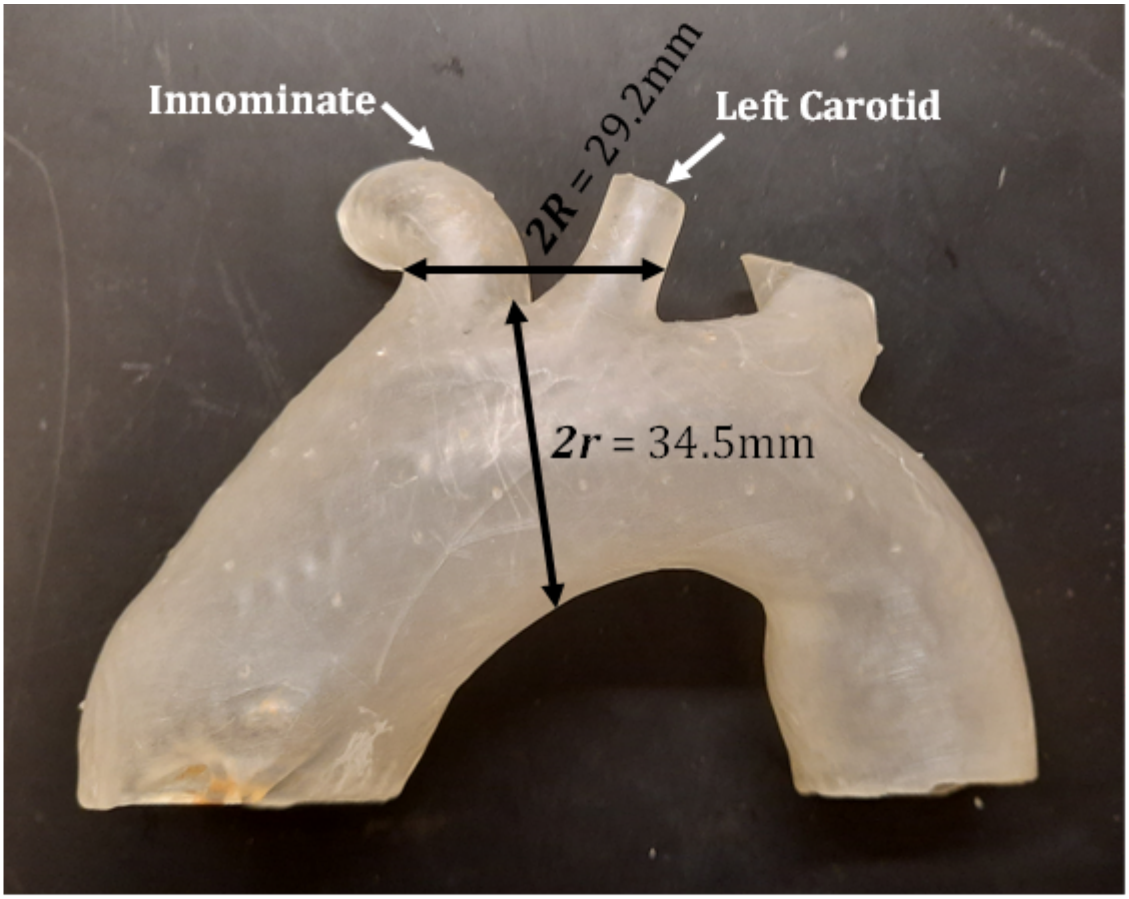


**Supplementary Figure 2.** Image of the rigid, 3D printed aortic arch model with key dimensions shown.
